# From initial RNA encoding to the Standard Genetic Code

**DOI:** 10.1101/2023.11.07.566042

**Authors:** Michael Yarus

## Abstract

Multiple experiments have shown that RNA binds chemically varied amino acids within specific oligoibonucleotide sequences. The smallest, simplest, and potentially most primitive RNA binding sites frequently contain conserved triplets corresponding to the Standard Genetic Code (SGC). Here, implications of such cognate coding triplets are calculated, combining them with an optimized kinetic model for SGC evolution. RNA-amino acid interactions at observed frequencies choose an SGC-like code, and, using the same mechanism, effectively resist alternative triplet assignments. Resistance to other kinds of coding is evident across varied code initiation scenarios. RNA-mediated assignments at experimental frequencies are sufficient to guide the ‘ribonucleopeotide transition’ (RNPT) to a modern code. This can account for extreme selection of the SGC among its astronomical code possibilities; very SGC-like codes are ca. 1/50 to 1/5 of codes within such a population. Nevertheless, full accounting depends on RNA affinities yet unmeasured. Such a code begins as mostly stereochemical, excludes mismatched assignments, and critically relies on properties characteristic of fusible microbes. After its RNPT in a partially assigned code, evolution accelerates definitively. Other assignment methods (adaptation, co-evolution, revised stereochemistry, LGT) likely complete the modern SGC because stable cellular intermediates with > 1 code exist, allowing compartmental code exchanges. Though initiated using chemical affinities, the 83 order-of-magnitude focus required to find a near-complete SGC among all possible codes was made by sequential evolutionary anthologies, in successive biological settings.

## Introduction

### An extraordinary bottleneck

Allowing unassigned functions, there are 1.8 x 10^83^ possible genetic codes with 20 functions and 64 triplets (1). Moreover, available primordial amino acids (2) greatly exceed selected biological forms (3). Carbonaceous meteorites contain at least 75 amino acids, a minority of which are biological (4). Further, it is clear that multiple bacterial codon assignments can be changed without severe biological consequences (5). Therefore, it is most striking that more than 250,000 codes in current bacterial and archaeal genomes (6) present only variants of the Standard Genetic Code using the same 22 amino acids (7).

Thus, modern biota descend from a single ancestral group. Early life on Earth went through an astronomically extreme bottleneck from which only one lineage, with its Standard Genetic Code, has emerged. Accordingly, much was determined by this event. Consider the choice of encoded amino acids (3, 8); some standard amino acids are inherited from a universal coding ancestor. More completely, the SGC embodies information from a crucial era hosting ancestors of all life on Earth.

### Initial coding

Inevitably, numerous amino acids were encoded before the advent of modern aminoacyl-RNA synthetase proteins (aaRS), which acylate RNA adaptors (present tRNAs) with specific amino acids for translation. This follows because aaRS are themselves complex polypeptides, with specific sites for at least three substrates, and specific active site geometries supporting catalyzed regiospecific aminoacyl transfer (9). Thus the aaRS era must have arisen from a simpler type of competent aminoacyl-RNA synthesis and translation, presumably mediated by RNAs (10–12). In particular, RNA-catalyzed amino acid activation (13) and aminoacyl transfer to RNA (11) have been reproduced and studied.

### RNA-amino acid interactions

Extensive experimentation shows that amino acids are bound specifically to small ribooligonucleotide sequences. Though originating with a natural *Tetrahymena* example, abundant further data exist (14–20).

In particular, RNA binding sites for amino acids have been characterized experimentally. Randomized RNAs were subjected to selection-amplification (21) after affinity chromatography for amino acids (22). Elution with free amino acids ensures that selected RNAs bind them. A summary of 464 such independently-derived sites for 8 chemically-varied amino acids, containing 7137 site nucleotides selected from 21938 initially randomized positions is shown in Table 1 (23). Local sequences within such binding sites contain an excess of cognate coding triplets for their amino acids, when compared to nearby non-binding-site sequences, which came from the same randomized RNA pools and passed through an identical purification. Interestingly, if the smallest binding sequences (the most likely primitive examples) are selected, inclusion of essential coding triplets is complete (24).

**Table 1.** Cognate codons and anticodons in RNA-amino acid binding sites. Description, references and details are found in (23). RNAs that bound to columns containing immobilized amino acids were isolated from randomized ribooucleotide sequences. Probabilities shown for abundance of coding triplets in amino acid binding sites were assessed by comparing originally randomized non-site RNA sequences with their associated binding site sequences, assuming that these abundances are equal.

|  |  |  |  |  |  |  |  |  |
| --- | --- | --- | --- | --- | --- | --- | --- | --- |
| probability | 3.9 x 10<sup>-5</sup> |  |  |  |  |  |  |  |
|  | UCG |  |  |  |  |  |  |  |
| probability | 4.0 x 10<sup>-26</sup> |  | 1.6 x 10<sup>-8</sup> | 3.2 x 10<sup>-131</sup> |  | 5.5 x 10<sup>-5</sup> | 2.7 x 10<sup>-13</sup> | 6.0 x 10<sup>-6</sup> |
| anticodons | CCU |  | GUG | UAU |  | GAA | CCA | AUA |
| *amino acids* | *Arg* | *Gln* | *His* | *Ile* | *Leu* | *Phe* | *Trp* | *Tyr* |
| codons | AGG |  |  | AUU |  |  |  |  |
| probability | 5.6 x 10<sup>-19</sup> |  |  | 8.0 x 10<sup>-110</sup> |  |  |  |  |

Every RNA site is independent because it has different sequences in initially randomized regions outside the binding site itself. In contrast, site sequences are repetitive, because amino acid binding must conserve atomic contacts with ribonucleotides. In fact, such sequence conservation is strong evidence that conserved coding triplets within binding sites serve essential functions. This is so because such triplets cannot be experimentally modified without disrupting binding function (25).

Table 1 links amino acids, their coding triplets and the probability of observed triplet concentrations in RNA binding site nucleotides. Probability cited is that cognate triplets are equally distributed between site and non-site sequences. A low probability threshold corrects for multiple trials (23): in Tab. 1, significant triplet concentrations have probabilities < 2.3 x 10^-4^.

Analysis of these RNA-amino acid binding sites has been questioned (26), but common objections are not compelling. Amino acid-ribonucleotide sequence associations do not rely on probability calculations. Site populations consist of highly repeated independent examples of one, or more, families of related site sequences (27). Cognate triplet sequences are directly observed as repeats among binding site nucleotides (also identified by biochemical criteria), so association cannot be a statistical artifact. Such data do not identify coding triplets as immediate contacts for the amino acids (an early idea tracing to coding in ‘holes’ on DNA: 28). But contact is not logically required, because conserved proximity of triplet sequences and cognate amino acids is sufficient to explain a later coding relationship. Moreover, it is sometimes asserted that the SGC mixture of primitive (e.g., Ile) and later amino acids (e.g., Arg) RNA sites is improbable, but it is argued below (Discussion) that this is expected. Similarly, seeking prebiotically plausible binding sites with few nucleotides commonly mandates moderate dissociation constants (23), but this is not a valid objection to site significance. Michaelis-Menten catalysis imposes rates dependent on forward 1^st^ order velocities, as well as affinities. Whole or fractional millimolar dissociation constants (23) are fully consistent with useful (bio)chemical velocities.

### Notable selectivity

Keeping in mind that Tab. 1 surveys only 40% of encoded amino acids, and that modern laboratory conditions may not wholly reproduce the Earth of 4 gigayears ago, observed RNA-amino acid interactions nevertheless suggest coding implications.

**Generality.** Six of eight sufficiently-tested amino acids show potential stereochemical ancestry.

**Variety.** Varied side chains exhibit RNA stereochemical links (Tab. 1): aliphatic hydrophobic (Ile), potentially catalytic (His), polar, strong H-bonding (Arg), planar aromatic (Phe, Tyr, Trp).

**Specificity.** For example, RNA distinguishes similar aliphatic sidechains, isoleucine and leucine (Tab. 1). An L-valine site distinguishes chemically similar D-valine, L-Leucine, L-isoleucine, and L-norvaline (29). An RNA binding site selected for phenylalanine can accept other planar aromatics, or reject tyrosine and tryptophan (27).

**Triplets.** No RNA affinities are associated with amino acid codons alone. All six amino acids sites link amino acids to cognate anticodons, thereby resembling associations in modern translation (30, 31).

**RNA-amino acid interactions provide a unique code origin.** Above, the astronomically vast universe of possible codes is emphasized. Selection of one possibility of ≍ 10^83^ requires explanation.

Thus an effective, specific mechanism for bringing RNA sequences together with amino acids is required. Such a mechanism almost identifies itself. It is difficult to imagine encoding without a physico-chemical interaction inducing proximity between specific amino acids and cognate coding sequences.

Thus, stereochemical origins are unique for two reasons. First, because of the need for choice among an astronomical code variety. Second, to juxtapose reactants and so (at least) entropically (32) catalyze encoded joining reactions.

Frances Crick supposed that the SGC might be a “frozen accident”, and “in its extreme form”, coding assignments could all be due to chance (31). But the SGC is not random (23, 33). But once evolved, the code would be resistant to change, because of the necessity to preserve pre-existing essential gene products (34). However, Crick anticipated that some amino acids might be initially encoded stereochemically (31). The finding that six of eight amino acids, of varied types, are related to cognate triplets (Tab. 1) suggests that chemical determinism rather than undirected chance may have a large role.

It is also possible that the code is ‘adaptive’, in the sense that it was selected to minimize phenotypic error when a coding mistake is made (35). Adaptive code evolution implies pre-existing coding whose assignments will be guided toward minimal error. So, adaptation does not initiate coding, nor choose among ≍ 10^83^ possibilities.

Evolution of the code might also be shaped by ‘coevolution’ with amino acid synthesis, with late-evolving amino acids also encoded later (36, 37). Coevolution is a persuasive idea, but offers no way to initiate coding, implement molecular association, or to make astronomically specific choices.

Given that the only apparent way to realize effective coding is with specific stereochemical interaction (38, 39), a mechanistic relation (below) between chemical and other code-ordering mechanisms will clarify code evolution.

### Chemistry for a code

Presumably cognate codons, anticodons, or both proximal to an amino acid binding site can be counted as stereochemical links, useable in forming a later code based on binding (23, 40). By this criterion, 6 of 8 examined amino acids have such a link (Tab. 1). Arg, His, Ile, Phe, Trp and Phe show evidence of stereochemically-originated coding, while no such evidence appears for Gln or Leu (23). In Tab. 1, 6 of 8 amino acids still show a stereochemical link if only anticodons are taken as significant (41).

Thus nine cognate SGC triplets are significantly present in RNA binding sequences for the six amino acids. However, CCU 3’/AAG 3’ are complementary anticodon/codons for Arg, so imply only one stereochemical assignment. UAU 3’/AUU 3’ are also both cognate for Ile, but are not complementary and so are separate chemically-determined assignments. So: 6 of 8 amino acids are putatively related to 8 SGC triplets. We will call this set of assignments “Stereo” to concisely refer to Tab. 1 chemical data. While extrapolation is not unique, extrapolation to 20 amino acids would implicate (20/8) x 8 = 20 stereochemical triplet assignments for 20 SGC amino acids. Thus, on simple assumptions, ≍ (20/61) ≍ 1/3 of amino acid-encoding triplets in the near-universal genetic code arguably arose from interactions between amino acids and ribonucleotide sequences. Extensions of observed 8 stereochemical triplets to complete coding tables are called _resi1 through _resi5 below, for ***r***andom ***<u>e</u>***xtension of ***<u>s</u>***tereochemical ***i***nitiation (supplementary data).

### A likely mechanism for coding table evolution

The path of least selection provides the most probable evolutionary route for a favored characteristic (42). Least selection for genetic encoding utilizes fusion of separate partial codes (43). Such code fragments might have been selected to serve varied functions, so fusing codes have continually increasing biological abilities. Increasing functionality makes fusions easily selected. In addition, code fusions implement evolution in parallel, so yielding complete codes more quickly (43). Given a prominent role for code fusions (44), either homogeneous or heterogeneous fusions leading to the SGC appear possible (45). That is, fusions may join further-evolved descendants of one original code, or may join codes from independent origins.

### The general objective

Below, amino acid-RNA associations are set within the shortest, most accurate code evolution known, to study their joint outcomes.

## Results

### Early amino acid-RNA interactions favor similar later code expansion

In Figure 1, code evolution environments initiate very differently: with either a single SGC-like assignment, or with a complete stereochemical SGC complement, as in **Chemistry for a code** above. Initial codes are completed (evolve to >= 20 assigned functions) by adding single random SGC-like assignments with specified accuracy (having probability Prand of randomness, e.g., without relation to the SGC).

**Figure 1.**
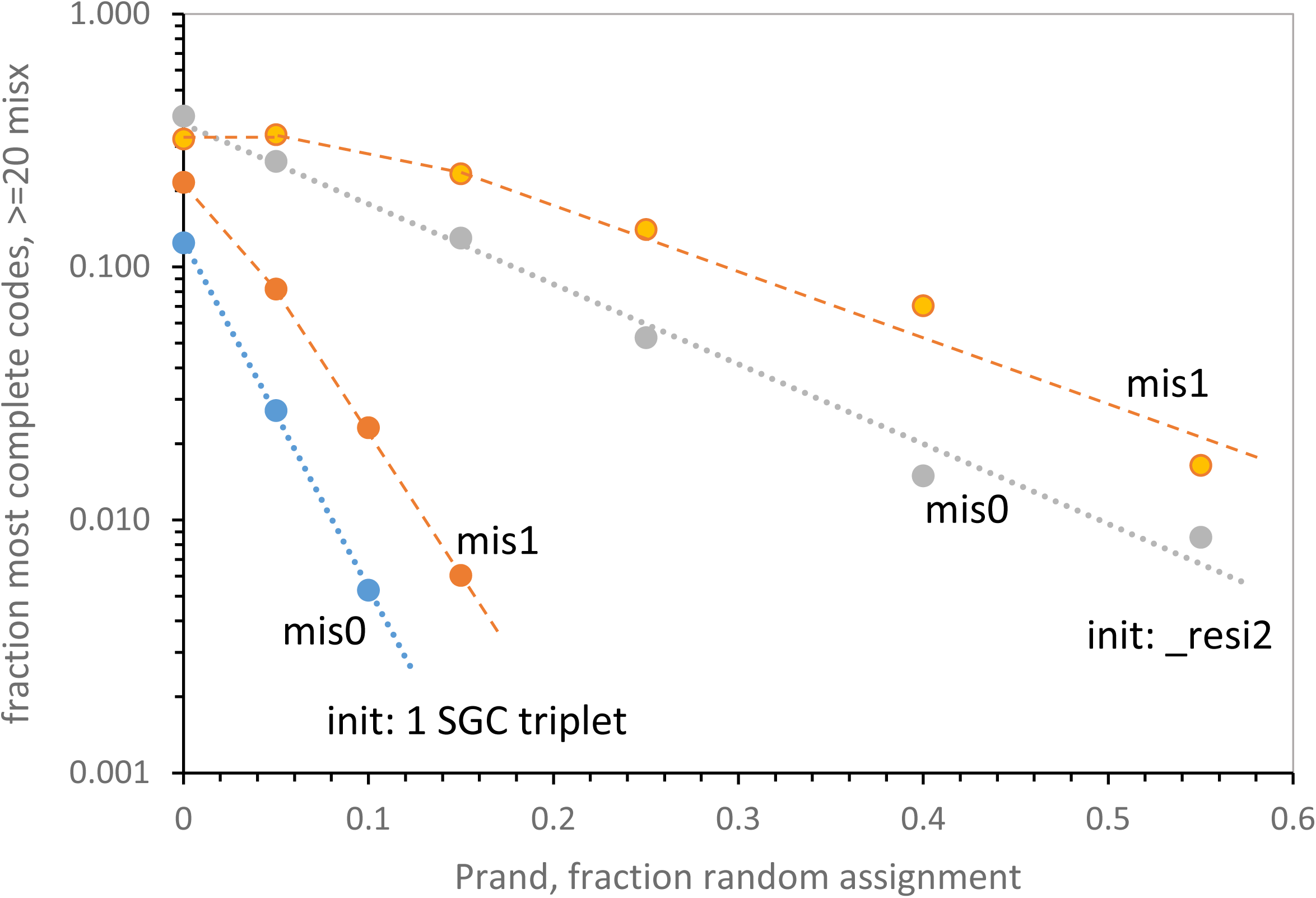
Coding table initiation alters tolerance for later assignments. Frequencies for near-complete codes (>= 20 assigned functions) with exact SGC assignments (mis0) and near-complete codes with one difference from SGC (mis1) versus the fraction of non-SGC (randomized, Prand) assignments throughout coding table evolution. The two pairs of curves compare evolution after code initiation with a single SGC triplet with initiation with evolution after a full complement of stereochemically-determined triplets (_resi #2; see supplementary information).

Fig. 1 plots frequencies of the most SGC-like final codes, having >= 20 encoded functions with no coding differences from the SGC (mis0) or with one non-SGC assignment (mis1). Upper Fig. 1 curves are for 20 initial stereochemically determined triplets, compared to lower curves for single-codon initial assignment.

Large differences result from distinct beginnings; initiation of code evolution with observed levels of stereochemical assignment makes completion of the code much more tolerant to error (randomness in later assignments, Prand). Thus, later assignments can contain errors, while still evolving to approach the SGC. Stereochemical initiation therefore gathers other compatible assignments to itself, allowing later divergence among assignments without making SGC-like coding inaccessibly rare. These effects are large; more-than-order-of-magnitude differences in final code abundances occur even for moderately greater random coding (Fig. 1).

### Amino acid-RNA interactions reduce acceptability of divergent assignments

Just above, initial SGC-like stereochemical assignment favors later similar SGC-like assignment. There is a complementary effect on non-SGC assignments, disfavoring infusion of later, conflicting assignments.

Fig. 2 compares incorporation from an entirely different coding scheme (ALT, supplementary data) into near-complete SGC codes, using three kinds of initial coding. Fig. 2A shows data for single random initiations, then continuation with single random SGC assignments. Fig. 2B shows initiation with the 8 experimentally implicated triplets of Table 1, then single random. Fig. 2C shows codes initiated with 20 stereochemically implied initiations appropriate to a complete coding table (**Chemistry for a code**, above), then single random. For each initiation, incorporation of an alternative code that has no assignments in common with the SGC (ALT) into the total population of codes (labeled **pop**) is compared to incorporation into the most complete code (labeled **code**).

**Figure 2.**
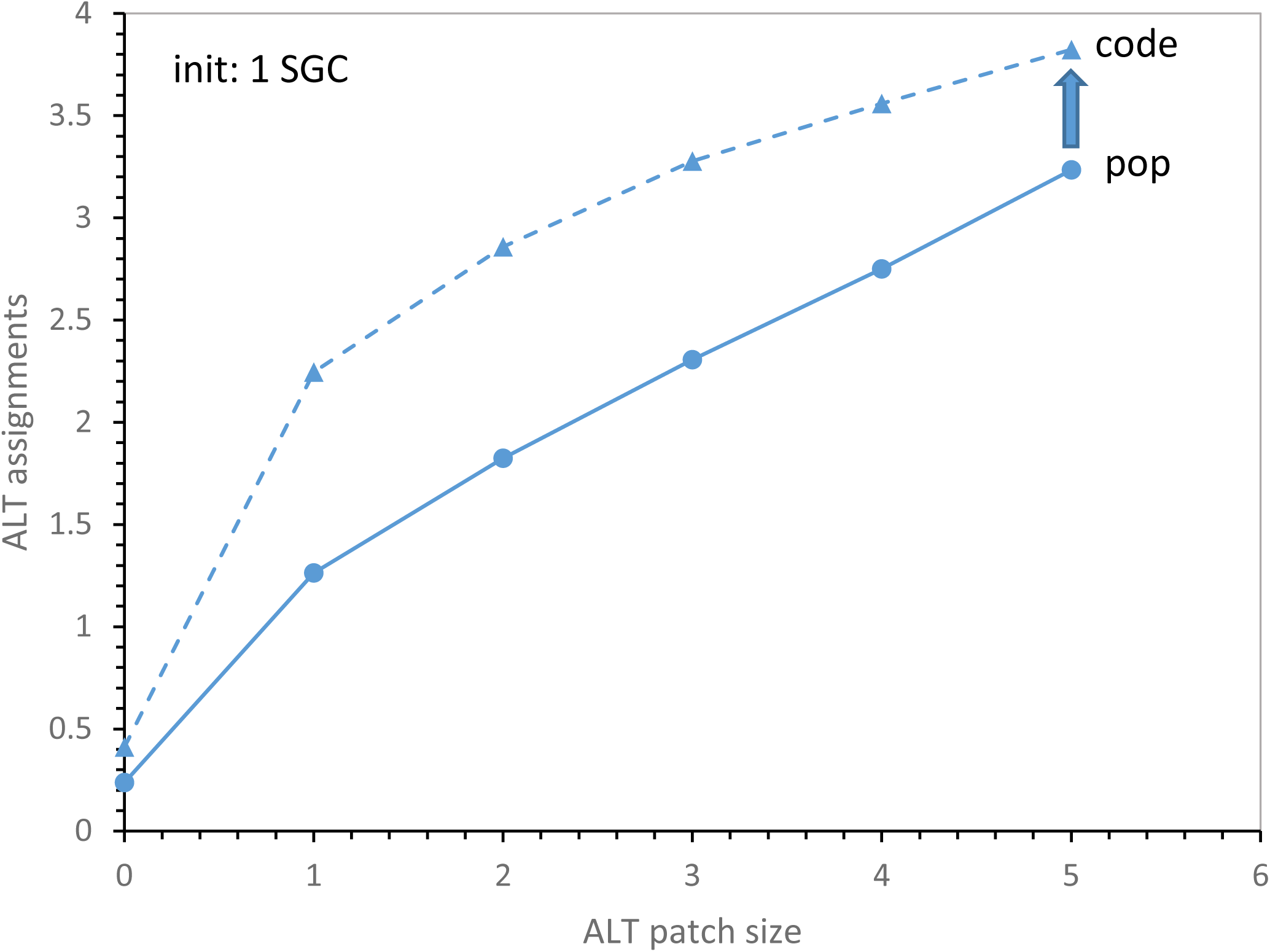
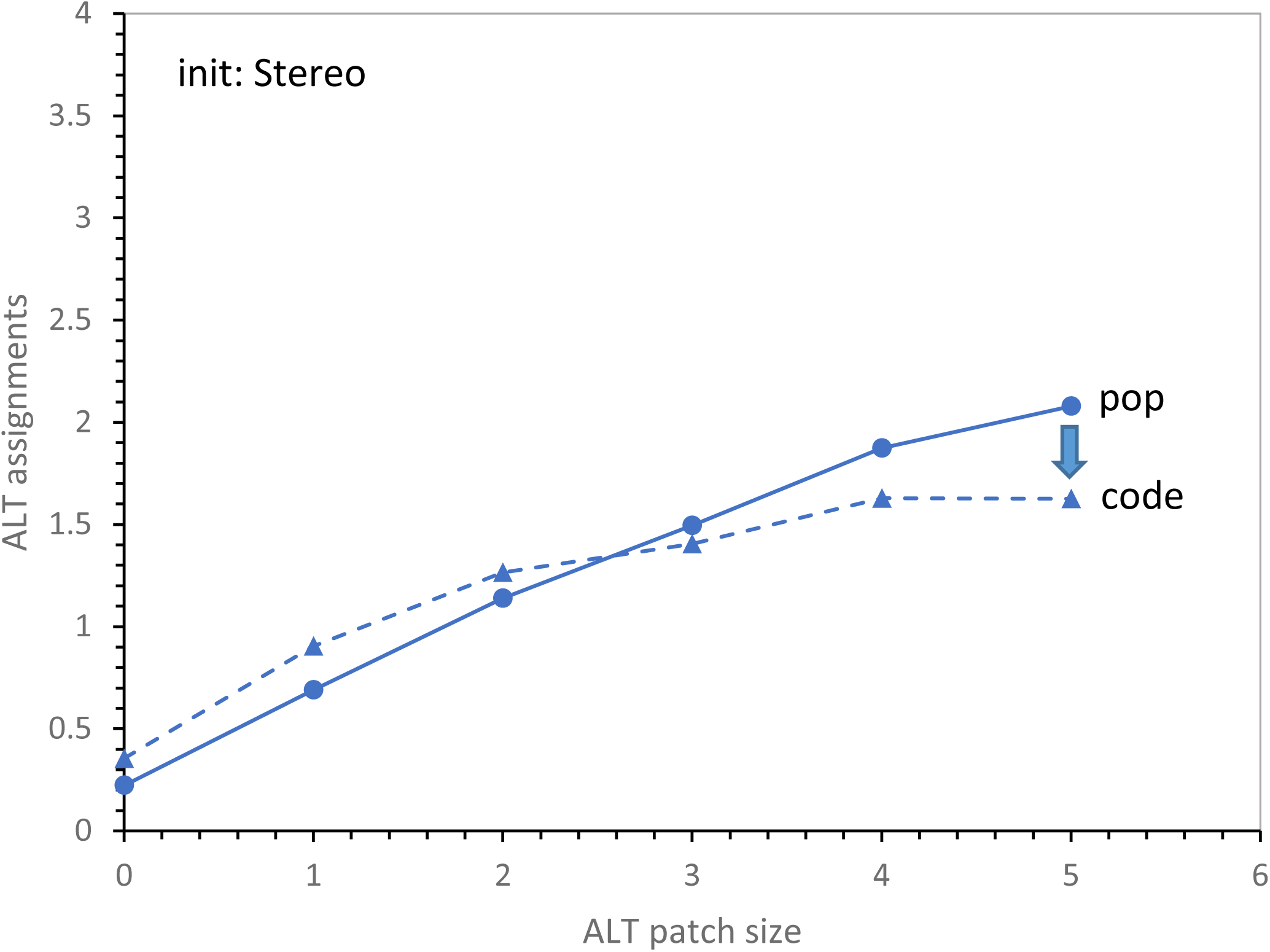
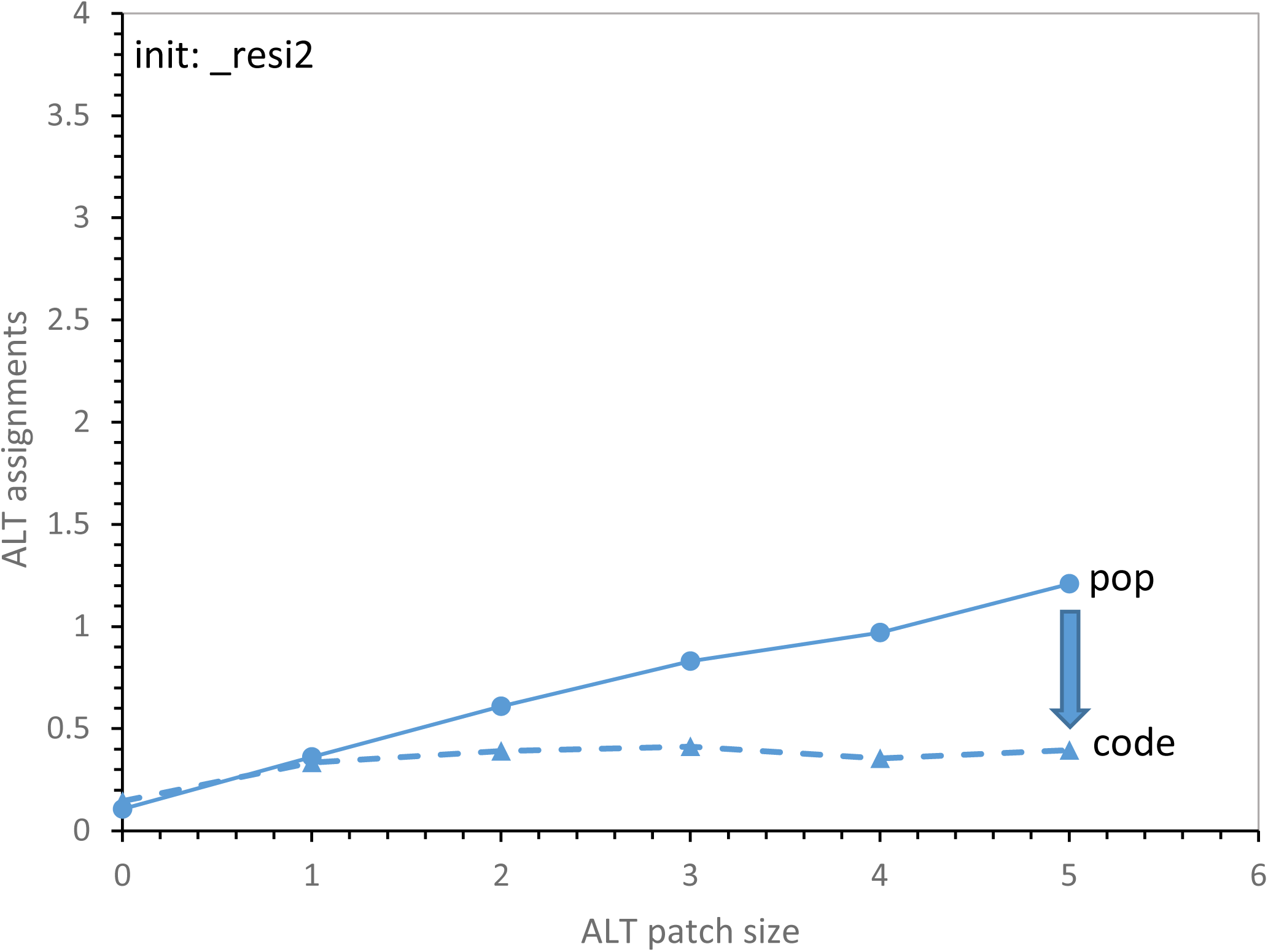
Coding table initiation alters acceptance of discordant coding assignments. A complete, full coding table having no assignments in common with the SGC (ALT, see supplementary information) was incorporated into evolving SGC-like codes, offering patch sizes of 1 to 5 randomly-chosen ALT triplets at a time. The number of accepted ALT-like assignments is plotted versus the number of triplets potentially transferred together. Each panel compares mean ALT-like assignment in the most complete codes (>= 20 encoded functions) with the mean in all evolving environmental coding tables at the time when the complete code arose. **Figure 2A – Code evolution initiated with a single SGC triplet.** **Figure 2B – Code evolution initiated with Table 1’s experimental triplets only.** See supplementary information for “Stereo” initiating assignments; 8 chemically-determined triplets. **Figure 2C – Code evolution initiated with a full chemically-determined triplet set, _resi2.** See supplementary information for _resi2; 20 chemically-determined triplets.

All panels share a Y-axis. Thus increasing initial stereochemistry, from single SGC assignments, to Stereo, to _resi2, decreases alternative coding (Fig. 2A, 2B, 2C). Though this has a simple basis (increasing numbers of codons are committed early), it is still a notable effect. For example, the average rightward code in Fig. 2A has seen 76 ALT assignments 5 at a time, but only 3.8 will be accepted in a superior SGC-like (>= 20 function mis 0) code.

Fig. 2 also distinguishes ALT incorporation into the entire code population (**pop**) from that into the most complete codes (**code**) for 1 ALT codon/assignment attempt to 5 codons per patch. Codes formed from single SGC triplets accept more alternative coding than exists in the general code population, thus using alternative assignments to complete their encoding (the arrow relates population mean to most complete codes, Fig. 2A). Coding restricted to experimental amino acid-triplet association (Fig. 2B) is intermediate, actually crossing over at 3 triplet patches. These intermediate initiations finally reject alien assignments, yielding most complete codes with fewer foreign codons (downward arrow) than the code population. Finally, initial assignment with stereochemistry extrapolated for total codes ((23), Table 1, Fig. 2C) discriminates strongly against alternative assignments, with fewer alternative encodings in most-complete codes at all ALT patch sizes (downward arrow, Fig. 2C).

So: alternative coding in a nascent SGC-like code is limited both with and without a body of early stereochemical assignments, but alternative encoding is particularly rejected with full stereochemical initiation, which accepts a mean of < 1 foreign assignment.

### Full stereochemical initiations differ greatly

For clarity, stereochemical initiations have been introduced as if they were uniform. But they emphatically are not. Code extension _resi2 (supplementary data) was used for illustrations (Fig. 1, 2), but other random extensions of known amino acid-RNA interactions have distinct outcomes. This is exemplified in Fig. 3A, which shows the mean time in passages required to evolve near-complete coding in 1500 evolutionary environments. When evolution of a stereochemically-initiated complete code requires more time, that code also admits proportionately more assignments from an independent coding system. Measurements show 8 triplets from evolved RNA binding sites (Stereo, Tab. 1, **Chemistry for a complete code)**, as well as 5 different extensions to completed codes (supplementary data). Time to complete a code varies 6.2-fold among stereochemical initiations, or 3.4–fold among the five completed initial triplet sets.

**Figure 3.**
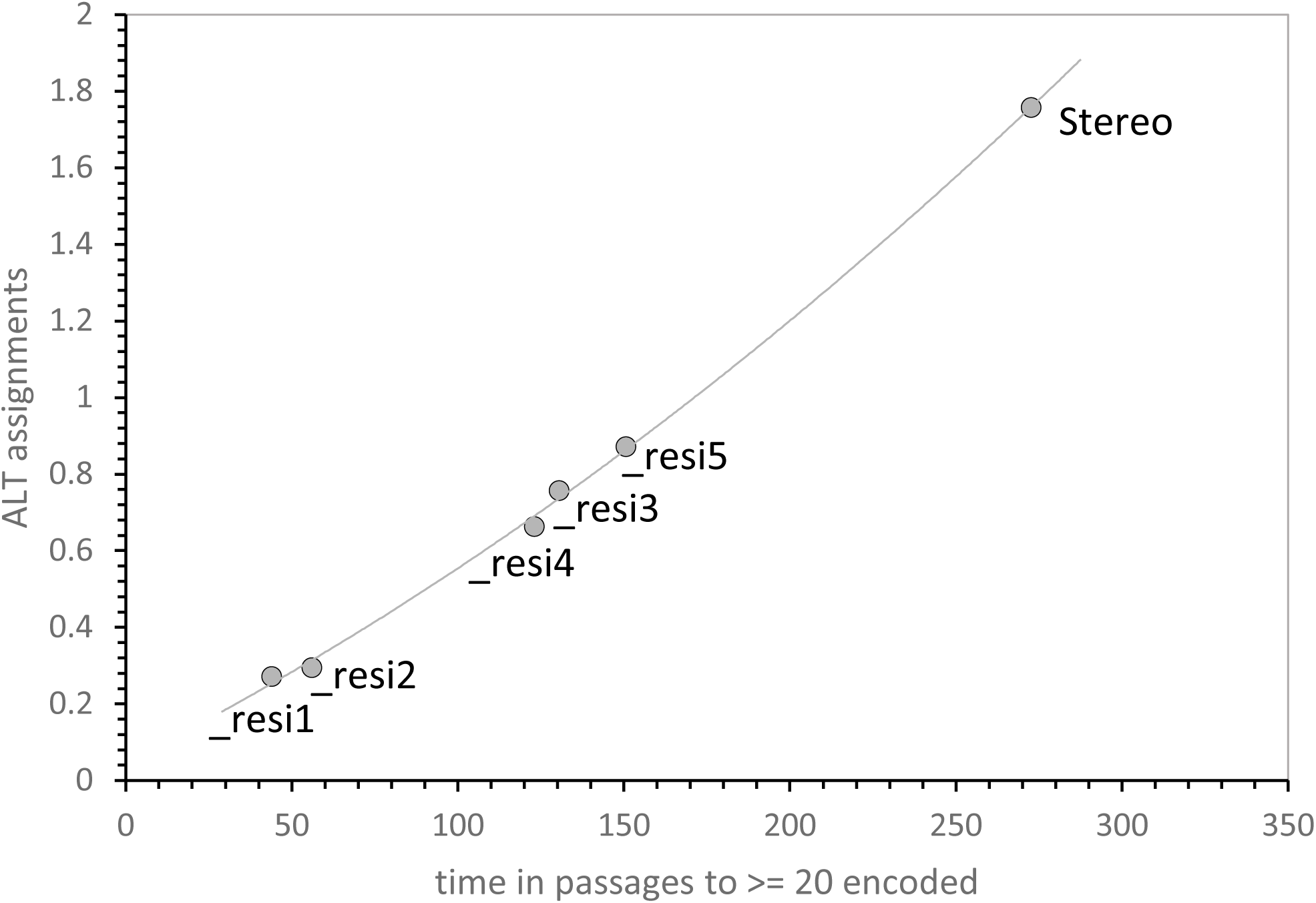
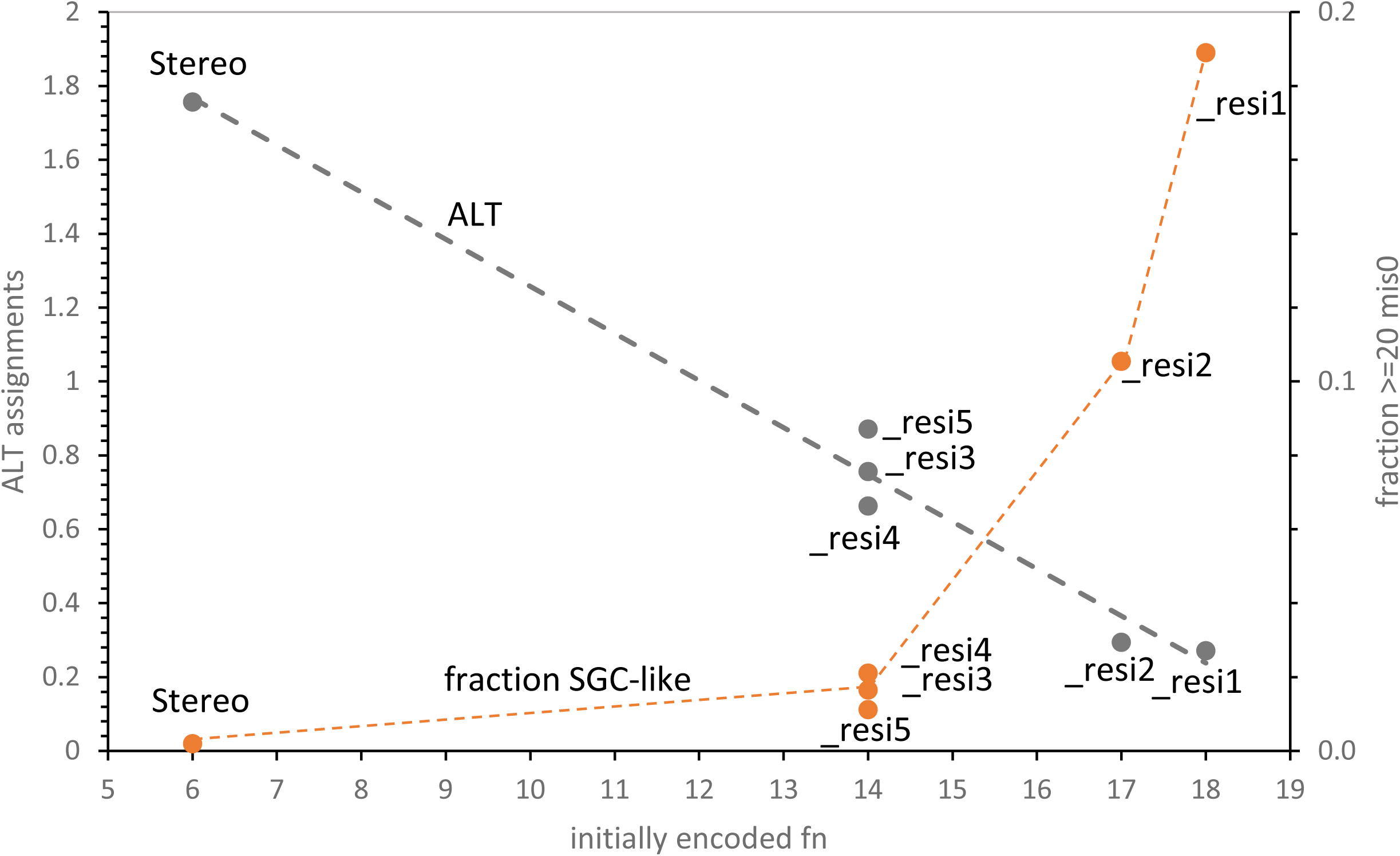
Code evolution is determined by the number of functions assigned by an initial coding set. The experimental triplet set (Table 1) surveys 8 functions (called ‘Stereo’). In contrast, _resi1, _resi2, _resi3, _resi4 and _resi5 are extended stereochemical triplet sets appropriate for complete coding tables (each with 20 determined triplets; see text **Chemistry for a code** and supplementary information). **Figure 3A – Incompatible coding assignments increase both time to evolve >= 20 functions and ALT triplet incorporation.** All codes evolved to encode >= 20 functions. **Figure 3B – Incompatible coding assignment decreases, and fraction of exact SGC coding increases, with the number of functions supplied by chemically-determined initiation.** All codes evolved to encode >= 20 functions.

### Stereochemical initiation differences have a simple rationale

Stereochemical initiations differ because of differing numbers of functions encoded. Time to evolve complete codes (Fig. 3A) and other evolutionary indices will vary with evolutionary distance to the SGC, in accord with a principle of least selection (42). Because full initial encodings are derived by adding randomly-chosen SGC triplets for non-studied amino acids to the 8 experimentally determined stereochemical triplets (Tab. 1), prospective full initiations based on the same observed 8 initial triplets differ in number of final functions encoded.

This is reflected in Fig. 3B, where the number of admitted ALT assignments is linearly related to the number of initiating encoded functions. The linearity extrapolates plausibly; with >= 20 functions initially (rightward in Fig. 3B), there is little or no room for alternative coding.

Such initial variation among stereochemical sets of triplets is yet more dramatic if a more informative distance to the SGC (45) is plotted. Such a distance measure reflects both amount of change and evolutionary accuracy. In Fig. 3B, the fraction of >= 20 function codes that are identical to the SGC (mis0 codes), starting with different initiations, is used. About 19% of all >= 20 function codes are identical to the SGC after _resi1 initiation (rightward, Fig. 3B). Observation of almost a fifth of near-complete codes identical to the SGC (Fig. 3B) concisely evokes the power of stereochemical initiation. It should be simple to select an SGC-like code among such an evolving code population. In particular, it appears that the challenge of a single SGC on Earth from 1.8 x 10^83^ potential coding tables can be readily met via selection from code fusion-division-late-wobble evolution, initiated with favorable stereochemical assignments. Alternatively: 19% of such nascent codes span the 83 order-of-magnitude SGC focus. Even for the least effective extensions, _resi3, _resi4 and _resi5, least SGC selection seeks ≍ 1 code in 50, still a seemingly short gap and simple task.

### Resisting inaccurate assignment

In Fig. 2 and Fig. 3, later code assignments are 15% random (Figure legends). These evolving code populations therefore benefit from Fig. 1’s effect. They find the SGC even when challenged by later departure from specific assignment, which would otherwise make an SGC inconveniently rare.

### Code populations reside in fusible microbes

Above results, in which the SGC is found despite somewhat unfavorable conditions, are due in part to exploitation of distribution fitness (1); selecting a favorable, less probable minority among a large, less-fit population. Fig. 2 populations range up to 10^5^ individual codes (Methods). Besides being numerous, they are able to fuse codes readily by fusing their coding compartments. Both conditions are readily rationalized if they are fusible unicompartmental microbes. Moreover, numerous microbes are also required to exploit the upper tail of distributed codes; that is, evolving by truncation, accepting only atypical best examples (42).

## Discussion

### Optimized evolution of coding tables

We draw the implications of observed stereochemistry using a Monte Carlo kinetic model of code evolution (1). The model has been optimized by determining the most rapid and accurate of 32 possible pathways for SGC evolution (45). Calculation constructs codes assuming that code evolution obeys standard rules of chemical kinetics (1). Different evolutionary pathways of can be compared using a principle of least selection (42), which asserts that the code most rapidly evolved and adhering most closely to known SGC assignments will require least selection; that is, will supply the most probable ancestors of the historical SGC.

Selected major conclusions of prior analysis are that only broadly validated evolutionary means are required to reach the SGC (45), that wobble was likely a later coding development, after most elementary assignments (1, 45, 46), that such wobble was more likely to have been simplified Crick wobble (45) than four-fold superwobble (47, 48), that the SGC was constructed by fusion of separately-evolved coding fragments (43), and that code division and re-fusion of evolved fragments was a major factor, notably enabling SGC evolution from one primary source of code assignments (45). Because fusing codes become unfit if they attribute ambiguous meanings to assigned triplets, fusion pathways purify coding, tending to converge on an underlying coding consensus, like the SGC (44). Moreover, fusion produces a prolonged elevation, or “crescendo” of changing codes, all very SGC-like. Within such a crescendo, SGC can be found by leisurely selection among as many competent codes as required (44, 45).

### Code fusion implies code convergence

Fig. 4A displays two merging code sources; if they use differing coding assignments, fused codes assign triplets ambiguously. This makes such codes less fit. Such ambiguous fusions become extinct. A minority of codes surviving fusion therefore has consistent assignments. That is, surviving codes converge to a consensus. This supplies the likely route to a coherent SGC (44). The same mechanism rejects completely independent codes (Fig. 2).

**Figure 4.**
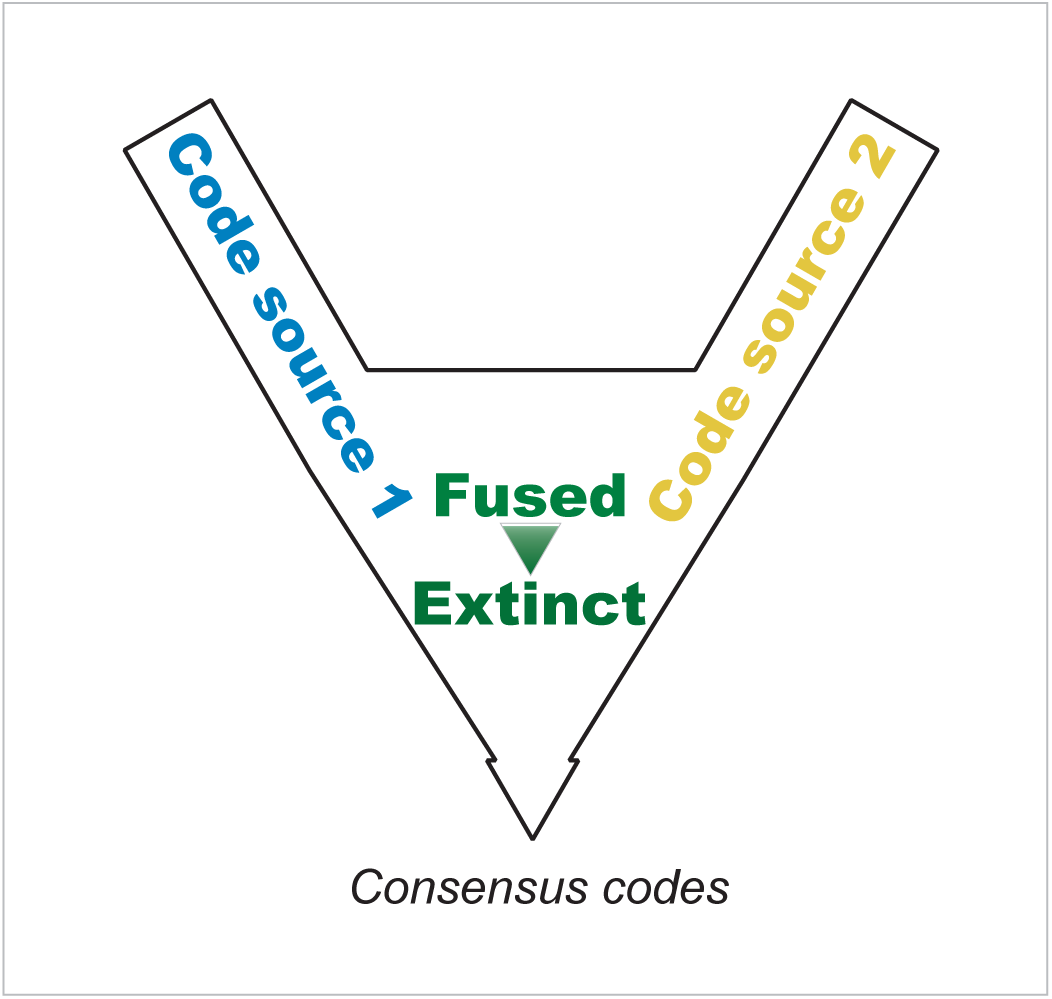
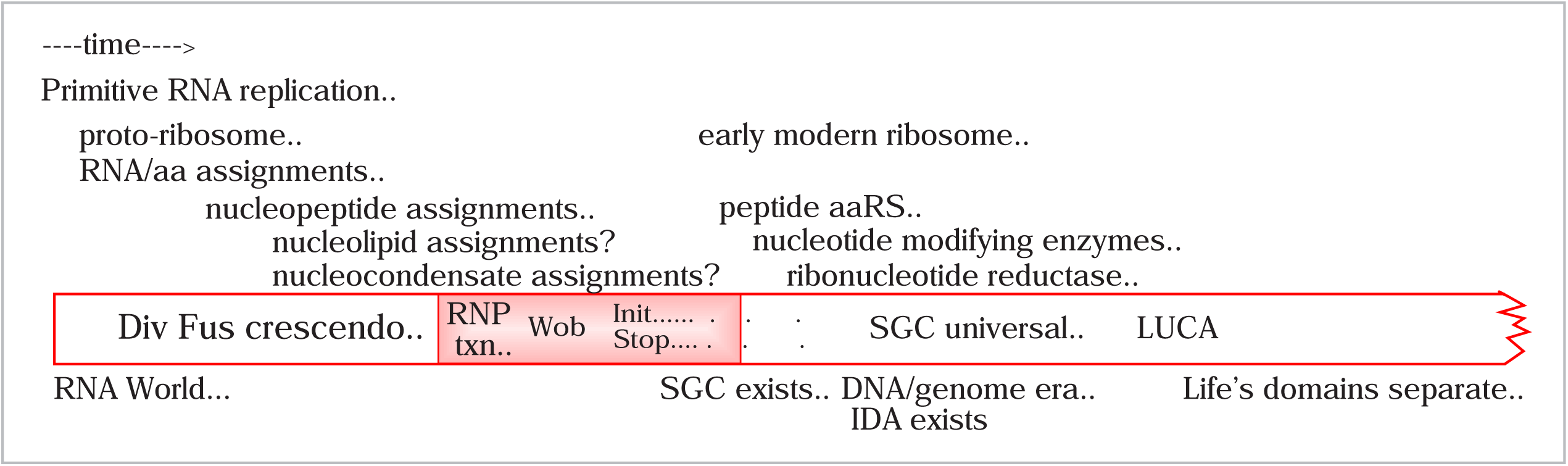
**Figure 4A – Unambiguous codes among fusion survivors converge to a consensus.** Code fusions that assign multiple meanings to individual triplets are less fit. Their extinction increases as codes become more complete, leaving a reduced number of more homogeneous codes for further evolution. **Figure 4B – An RNA era for the code.** As the only evident means of specific code initiation, chemical interactions between oligoribonucleotides and free amino acids define an early RNA coding era (see **RNA Coding in context**). Such coding is sufficiently complete to produce the RNPT, yielding highly competent structural and catalytic peptides. Stereochemical assignments tend to exclude independently-originated coding assignments, but these may be added to stereochemical precursor codes during later evolution.

Such discrimination is more strict for complete stereochemistry (Fig. 2C) than if restricted to experimentally confirmed effects (Fig. 2B), and is least exclusionary in the extreme case that random SGC codons are added at random times, one-by-one (Fig. 2A) instead of being added in an initial set (Fig. 2B, 2C). If SGC codons are added one-by-one, the most complete coding tables actually import a small number of alternative assignments to complete themselves (Fig. 2A). In contrast, initial full-code stereochemical assignments almost exclude alternative assignments, accepting less than one codon per near-complete code (Fig. 2C).

### But full stereochemical initiations differ

Completed sets of potential stereochemical assignments differ greatly among themselves (Fig. 3, supplementary data). These differences affect most properties of interest; but 3.4-fold in time to evolve near-complete codes, and 6.5-fold in the number of alternative assignments admitted (Fig. 3A). This is due to varied initial coverage of the SGC: differing sets of triplets with the same stereochemical assignment frequency nevertheless initially assign different fractions of total SGC functions. The evolutionary path from such initiations to the SGC therefore has different lengths, requires varied times (Fig. 3A), and allows different proportions of unlike assignment (Fig. 3B). This conclusion does not require extrapolation: even prior known stereochemistry limits alien assignment to less than 2 codons (Fig. 2B) of the 64 possible in full codes. A persuasive estimate of alternative coding within the SGC will require a completed stereochemical census (23) for SGC triplets.

### Magnitude of the RNPT

The RNPT solves an acute problem. Early coding must be substantially complete in order to encode initial peptide aaRS, peptide factors, and other proteins required for near-modern protein synthesis. Fortunately, minimal amino acid complexity required for functional proteins has been extensively studied. Tertiary folding is the simpler requirement, and a 3-fold symmetric 140-aa fibroblast growth factor folds well when composed of 12 or 13 amino acids (50), rather than the standard complement.

Enzymatic folded forms are a more complex and relevant requirement for the current argument. Chorismate mutase, a 93-amino acid enzyme with a straightforward alpha-helical structure, has in vivo activity if reduced to only 9 amino acids (51). A 213-aa orotidine phosphotransferase (from PRPP) is active if reduced to 13 kinds of amino acids (52). A reconstructed ancestral bacterial NDP kinase functions with excellent thermal stability and up to 1% control activity with 13 or 14 amino acid alphabets (53). A protein of particular relevance, small ribosomal subunit protein uS8, with 130-aa, binds its RNA if composed of 15, 14 or 13 amino acids (54). Notably, though less stable than true thermophilic homologues, reduced uS8 proteins still have high stabilities, half-melted at ∼85°C.

In short, 9 to 15 different encoded amino acids can construct structurally and chemically functional peptides hundreds of amino acids long, with a frequent threshold at 13 different amino acids. The extrapolated stereochemical encoded complement of 15 amino acids (**Chemistry for a code**, above) therefore suggests that RNA can encode the initial RNPT.

Thus an initial code relying on RNA chemistry alone probably encoded polypeptides with major structural and enzymatic capabilities (Fig. 4B). After the resulting RNPT, evolution to modern translation and the full SGC is enabled, requiring exquisite adaptation between proteins and RNAs (55) and wide-ranging ribonucleotide refinement for coding (56, 57), but not radical evolutionary innovation.

### Life’s tempo and mode after the RNPT

A characteristic of RNA catalysis is that, though certainly more versatile and faster than once imagined, it is still limited and slower, compared to protein catalysis (58). Biocatalysis will accelerate by orders of magnitude after the RNPT brings peptide groups to bear on (bio)chemical transition states. The addition of histidine to coding (59, 60) and peptides (61) will likely, even alone, have a notable catalytic impact. As a result, cell activities, particularly including cell division, accelerate through the RNPT. Evolutionary success is almost synonymous with faster growth (62); thus the definitive early code plausibly emerges from post-RNPT cell divisions. RNPT transitioning cells race away from those with slower, less versatile catalysis, and we are their inheritors.

### DNA appears after the RNPT

Deoxyribose is not among easily accessible 5-carbon sugars (63) and the unusual radical catalysis universally required to generate it from ribose (64) would require biosynthesis of a sophisticated catalyst. This indicates that the DNA genome appears substantially after the RNPT, as shown in Fig. 4B.

In fact, consolidation of the DNA genome and thus Darwinian descent itself might have depended on pre-existing peptide encoding/translation (42) and thus on the RNPT. Effective post-RNPT translation of numerous encoded peptides sharply elevated the benefits of accurate inheritance.

### Fate of early assignments after the RNPT

Crick’s intuition about the ‘freezing’ of the code (31) applies to the RNPT (Fig. 4). Even with dramatic changes attending the appearance of peptidyl aaRS and ribosomes, RNA assignments would likely be mostly preserved in the early RNPT era to continue synthesis of critical peptides (65). So early triplet assignments likely survive in the SGC, though the machinery selecting them has changed. Separating RNA-era assignments and more complex RNPT assignments is clearly worth specific thought.

For example, amino acids assigned multiple sets of codons may have been assigned at two times, in two different ways. Accordingly, positing that Table 1 presents RNA-era encodings is tempting. But this is <u>not</u> plausible; the truth must be more intricate. For example, arginine has two sets of codons - CGN and AGR, has a complex biosynthesis (66), probably was encoded later (67). Arg exhibits multiple affinities to cognate RNA sequences (14, 15, 23, 68). Some Arg assignments can reflect RNA stereochemistry used for assignment after Arg biosynthesis became efficient in the RNPT-era, while other assignments reflect multi-molecular affinities, possible only in the RNPT-era. Accordingly, even complex amino acids, with late biosynthesis, can be encoded via cognate RNA affinity. However, genuinely persistent RNA-era assignments can exist (consider Ile, Tab. 1), as well as assignments requiring RNPT molecular collaborations and biosynthesis. More study may disentangle possible encoding histories.

### The SGC probably does not have a single origin

Present calculations therefore imply that there was an initial era almost entirely stereochemical, then a second era permitting other kinds of amino acid/triplet assignment. It thereby explains why a metabolic code origin (36) cannot be complete (69, 70) why a completely adaptive origin (35) becomes difficult to rationalize (71), why a random origin is improbable (72), and indeed, why even a stereochemical origin (23) will not explain every coding assignment (73).

### Coding is partial at the RNPT

Above, coding is incomplete at the initial RNPT, likely biased toward early and RNA-interacting amino acids. How is the SGC completed?

### Code logic can be more accepting after the RNPT

Abstractly, RNPT coding might continue to reject foreign assignments via fusion incompatibility. But an appealing hypothesis is: as the biome multiplied, endocytotic cellular fusions became more probable. Differing codes would coexist in one cell, in different membrane-bound compartments. Such persistent proximity between divergent codes, post-RNPT initiation, can elevate the frequency of successful code fusion via LGT.

Somewhat similar later events are well-studied. During the DNA era (Fig. 4B), massive multi-directional genetic transfer between compartments was the rule. For example, extensive LGT occurred from the alpha-proteobacterial precursor of the mitochondrion to the protoeukarotic nucleus (74). 10-16% of the modern mitochondrial proteome appears to persist from its bacterial ancestor, and at least 842 different initial protobacterial gene families have been identified in other modern cellular locations, including the cell nucleus (75). Gene transfer between DNA compartments was evidently strongly favored. This could augment the code, as well as promoting other evolutionary joinings.

### The RNPT in coding history

Fig. 4B puts the RNA era in historical context, specifically respecting the intricate history of genetic coding. First letters of the figure’s text entries mark time-of-origin for various events.

From left to right: an RNA world requires the ability to replicate RNAs accurately, though eased by the finding that active translational RNAs can be as small as tetramers, pentamers and hexamers (76–78). Initial coding assignments are made using RNA affinities alone (23, 40). Short, simple encoded peptides can be made on an RNA proto-ribosome, before a fully functional RNPT ribosome exists (79). As peptides proliferate, they can join with ribonucleotides to add to the specificity of amino acid binding and triplet assignment (80, 81). The possibility that other ancient molecules also joined RNA to form later amino acid binding sites should not be neglected: polar lipids (82–85) and condensates (86) mediate other well-characterized RNA associations.

Early assembly of the code relies on the code division-fusion crescendo which helps the code converge to the SGC, also speeding its evolution by allowing parallel progress and presenting a prolonged series of close SGC relatives, facilitating the code’s optimal least selection (43–45). A set of assignments results, more-than-sufficiently complete to make the RNPT (shaded region, Fig. 4B) to complete coding. RNA-peptide collaboration in coding functions appears during the RNPT, yielding wobble (46) and initiation and termination. Translation initiation (87) and termination (88) change during both early and late SGC evolution, evolving over a prolonged period that continues after the separation of major biological domains. Wobble is a late event during triplet assignment, because accurate wobble requires a large allosteric ribosome (89) and a refined tRNA structure (90, 91). Additionally, early wobble specifically and greatly hinders code completion (46).

After the RNPT begins, accurate coding of substantial peptides enables the appearance of an early but recognizably modern ribosome, usually modeled as the predecessor of the modern peptidyl transferase region (92, 93). Near-modern coding capabilities produce the protein aaRS (94), and deoxyribonucleotides (64). Thus, DNA, the genome and Darwinian evolution arise. Accordingly, the Initial Darwinian Ancestor emerges (95). These events are accompanied by nucleotide modifying enzymes that refine the codon-anticodon complex (56, 57). One complete code becomes the universal SGC (96) and diverges from LUCA to produce a modern multi-domain biota employing the Standard Genetic Code (97).

### Biology as Anthology

This adage defines life via its ability to combine advantages evolved separately (45). Here a prior ambiguity is resolved (45): efficient code division-and-fusion SGC evolution defines a unified route to the final code, but also seems to offer code fusion from varied origins. Fusion into a nascent SGC is possible but limited, decreasing as chemically-based initial assignment increases (Fig. 1, 2, 3). Thus, code fusion suggests, but also tenaciously limits, truly foreign code joining (44).

The historical SGC is thus a constrained code anthology. It still bears evidence of early stereochemical sources, conserved because assignments must be substantially conserved in the RNPT to preserve prior peptide capabilities. But early non-stereochemical RNA-era patches are small or nonexistent, dependent on the precise stereochemical history of the code (Fig. 2, 3, 4). Later, non-stereochemical contributions remain to be resolved, but this can be clarified by practical experiments.

### Biology as Anthology; sources of a 10^83^-fold convergence

In the RNA world, code fusion combining RNAs making specific chemical contacts with amino acids defined, restricted and homogenized a partial, but competent, code, possibly descended from a single ancestor (45). The most complete fused RNA codes supported a decisive RNPT, which speedily established its superiority. Then, in the RNPT era, a different code fusion mechanism possibly added to coding, adapting it to more competent, complex cells. In the DNA era, accurate, near-complete codes competed: a broadly-adapted victor became the SGC (96), efficiently encoding LUCA’s diverse proteins (98). The SGC is the product of three sequential anthologies: first joining the reactions of RNAs (23), then joining the competence of RNA and peptides (43), then joining proficiencies of populations (96).

## Methods

Monte Carlo kinetic calculations (1, 45) were performed in console mode within the Pascal development environment Lazarus 2.2.ORC1 using the FPC compiler 3.2.2, hosted on a Dell computer running 64-bit windows 10 Enterprise, v. 22H2, with an Intel i9-8950HK CPU @ 2.90GHz and 32 GB of RAM.

**Figure 1 results - Effect of differing initiations on later code acceptability.**

1500 evolutionary environments ultimately containing ≍ 63,000 to 72,000 total coding tables were surveyed after initiation with a single SGC codon, and evolved until a code with >= 20 assigned functions appeared in the environment. Assignments after the first had probability Prand of being random, thus were not constrained to be SGC-like. Pmut=0.00975, Pdec=0.00975, Pinit=0.150, Pfus=0.00100, Ptab=0.080, Pwob=0.0000, Pdivn=0.500. Pmut is the probability/passage of a related assignment to an unocupied neighbor codon related by a single nucleotide change, Pdec is the probability/passage of selected codon assignment decay, Pinit is the probability/passage of initial assignment to a new coding table, Pfus is the (probability/passage-coexisting table) that randomly chosen codes will fuse, Ptab is the probability/passage of initiation of a new coding table, Pwob is the probability/passage of the initiation of wobble coding (always 0.0, late coding, here), Pdivn is the probability/passage that a random existing coding table will divide identically (1, 45).

1500 environments contained ≍ 35,000 to 48,000 tables initiated with full stereochemical _resi2 (see supplementary information), then evolved until a code with >= 20 assigned functions appeared in the environment. Assignments after the initial set had probability Prand of being random, thus were not constrained to be SGC-like.

**Figure 2 results - Differing initiations restrict acceptance of alternative assignments.**

**Figure 2A – Codes Initiate evolution with one SGC codon** - 1500 evolutionary environments ultimately containing ≍ 96,000 to 109,000 total coding tables were evolved to yield a code with >= 20 assigned functions, as in Figure 1, except that with probability Ptab per passage, a group of 1 to 5 assignments could come, randomly, from coding table ALT (supplementary information) which has no assignments in common with the SGC. Pmut=0.00975, Pdec=0.00975, Pinit=0.150, Pfus=0.00100, Ptab=0.040, Pwob=0.0000, Pdivn=0.500. In addition, evolution included random assignments at Prand = 0.15.

**Figure 2B – Codes Initiate evolution with 8 experimental stereochemical triplets only (Stereo; Table 1)** - 1500 evolutionary environments ultimately containing ≍ 69,000 to 90,000 total coding tables were evolved to yield a code with >= 20 assigned functions, as in Figure 2A, except that with probability Ptab per passage, a group of 1 to 5 assignments could come, randomly, from coding table ALT (supplementary information) which has no assignments in common with the SGC. In addition, evolution included random assignments at Prand = 0.15.

**Figure 2C – Codes Initiate evolution with 20 stereochemical triplets (**see text: **Chemistry for a code)** - 1500 evolutionary environments ultimately containing ≍ 32,000 to 37,000 total coding tables were initiated with _resi2 and evolved to yield a code with >= 20 assigned functions, as in Figure 2A, except that with probability Ptab per passage, a group of 1 to 5 assignments could come, randomly, from coding table ALT (supplementary information) which has no assignments in common with the SGC. In addition, evolution included random assignments at Prand = 0.15.

**Figure 3 results - Complete stereochemical initiations differ markedly in final code properties**

Completed stereochemical initiations _resi1 through _resi5 (20 stereochemically-determined triplets: described in **Chemistry for a code**) are shown in supplementary data. Probabilities are as in Fig. 2A.

**Figure 3A – Time to evolve and ALT assignments for different initiations are related.** Coding tables evolved to >= 20 assignments with possible ALT assignments and initiation with experimental stereochemical assignments from Table 1 (‘Stereo’) or estimated full-coding-table assignments _resi1, _resi2, _resi3, _resi4 and _resi5 (text **Chemistry for a code**; see also supplementary information). Other conditions as in Figure 2A.

**Figure 3B – Stereochemical initiations differ because they differ in number of initially encoded functions.** ALT assignments accepted and the fraction of very SGC-like codes (>= 20 assigned functions and entirely SGC assignments) are plotted versus number of stereochemical functions initially encoded. Other conditions as in Figure 2A.

---

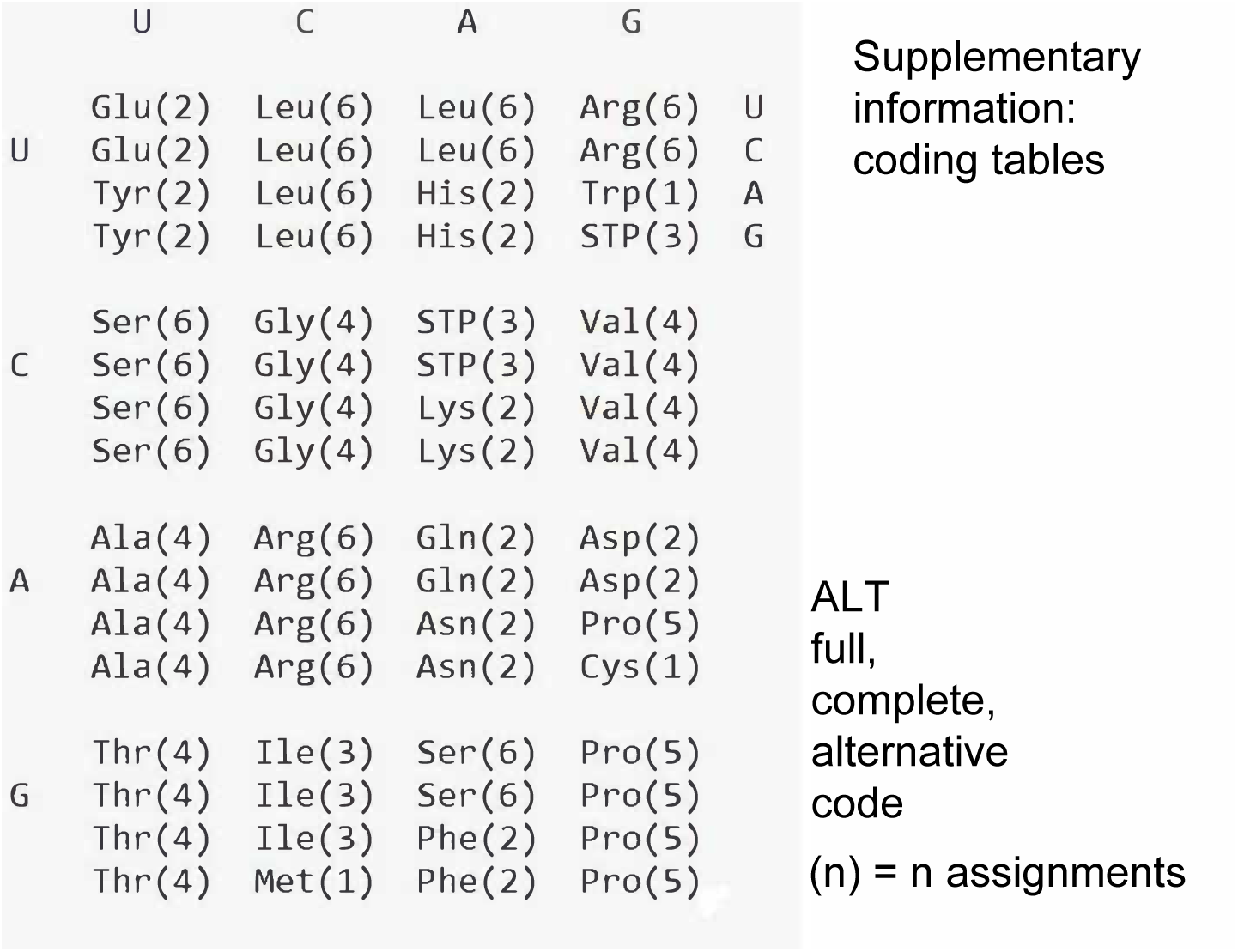
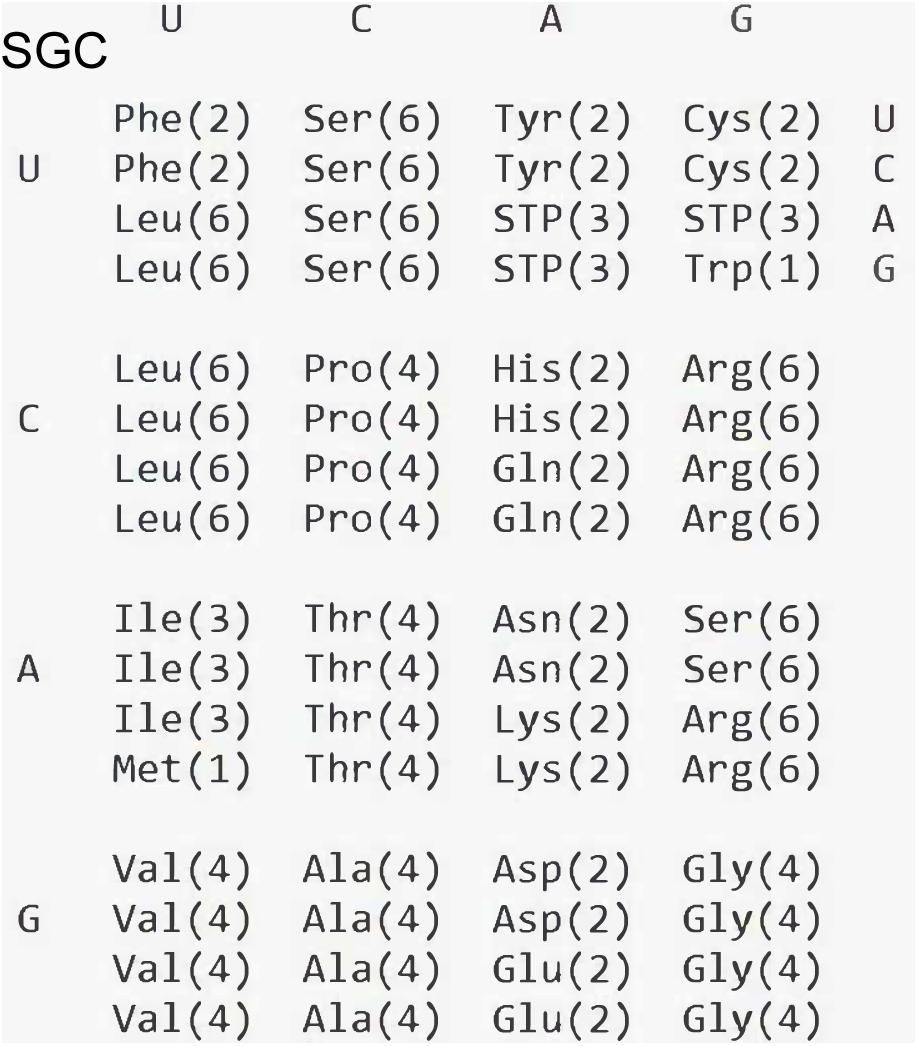
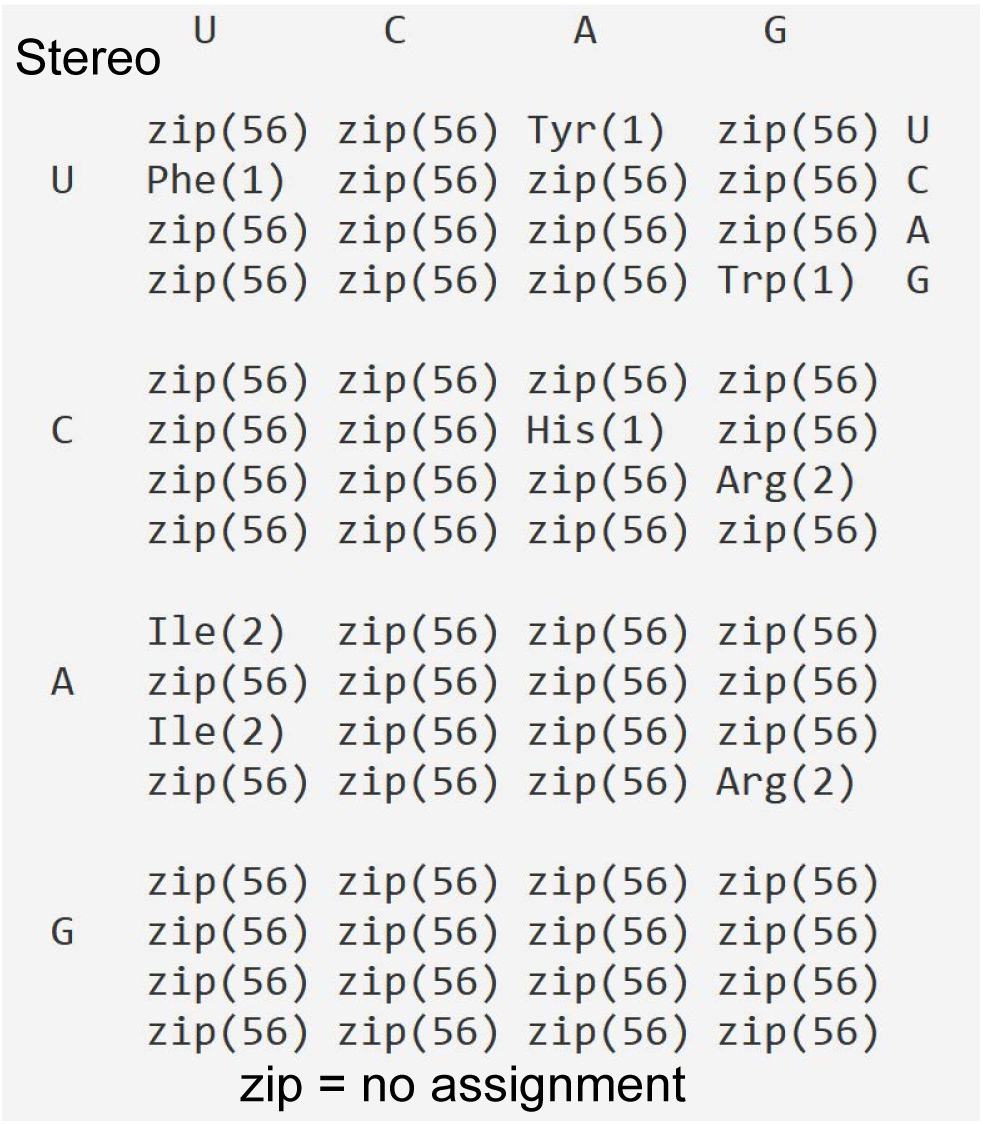
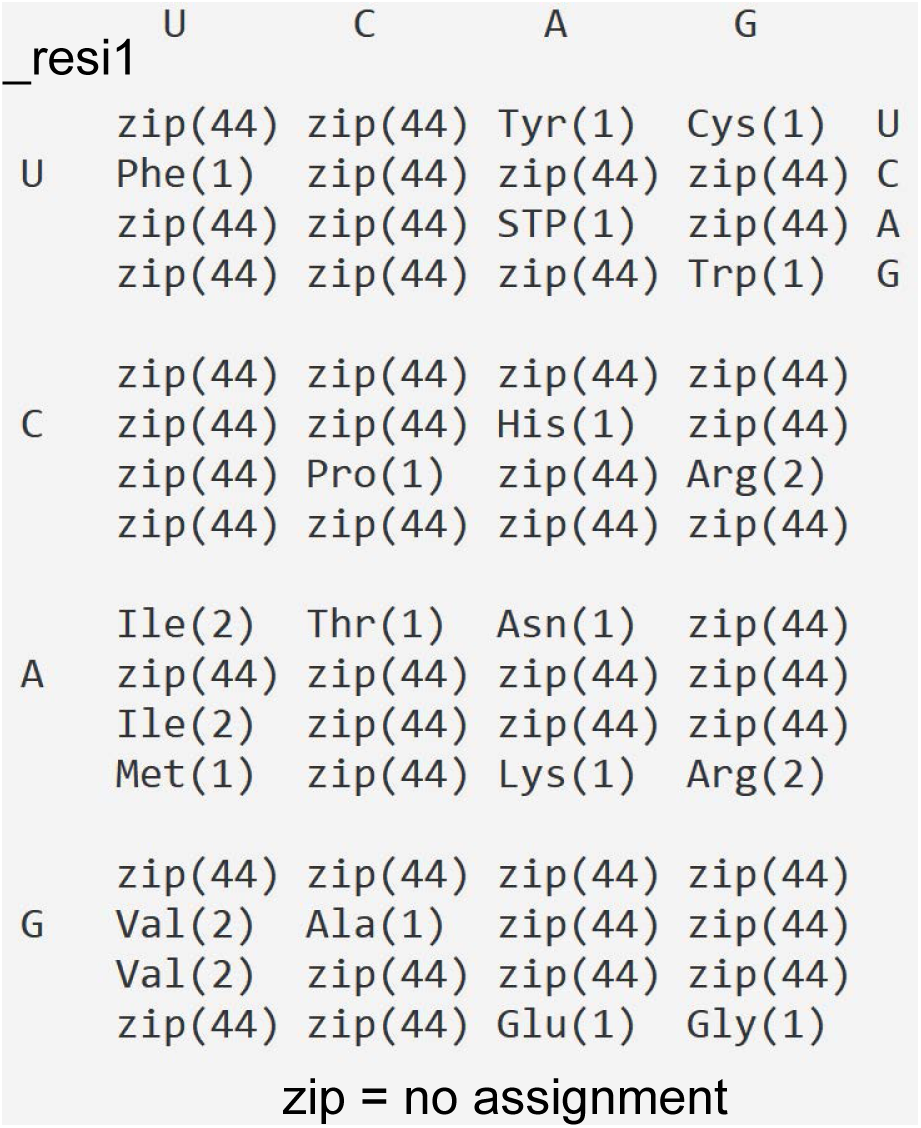
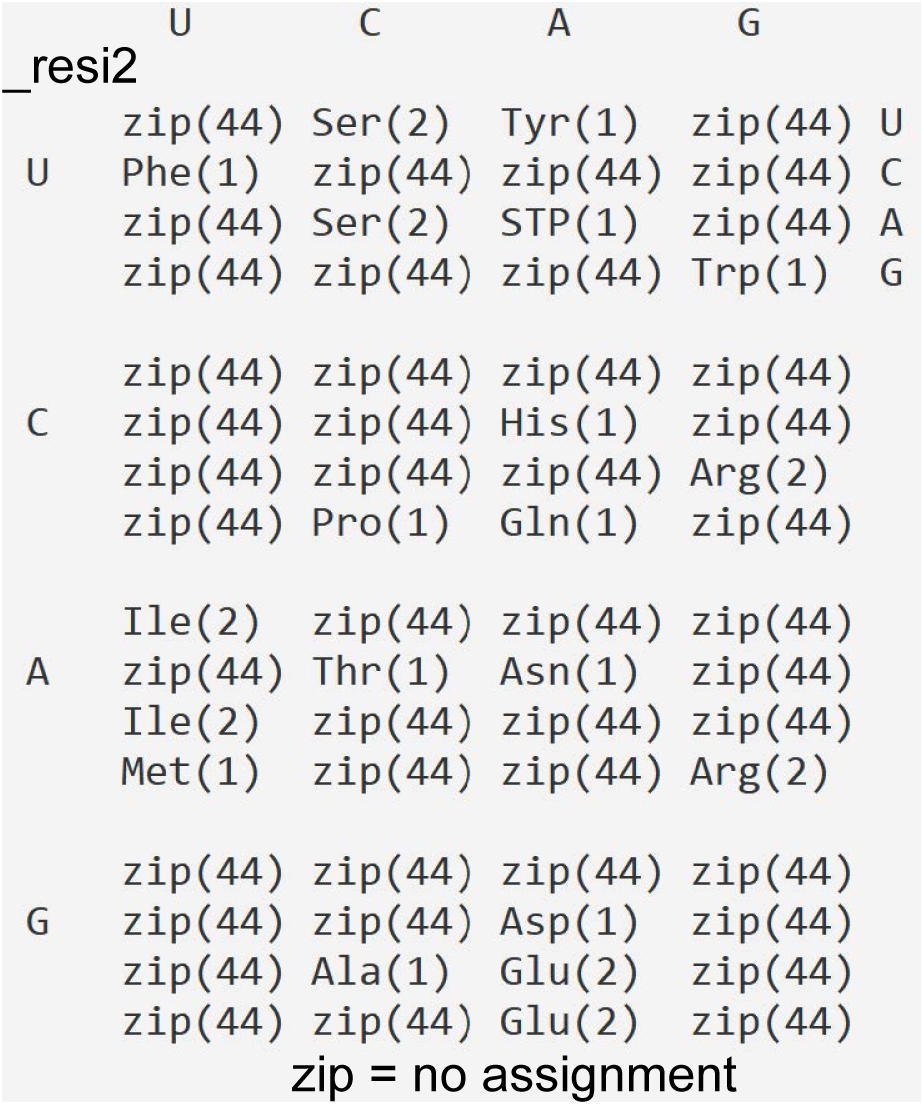
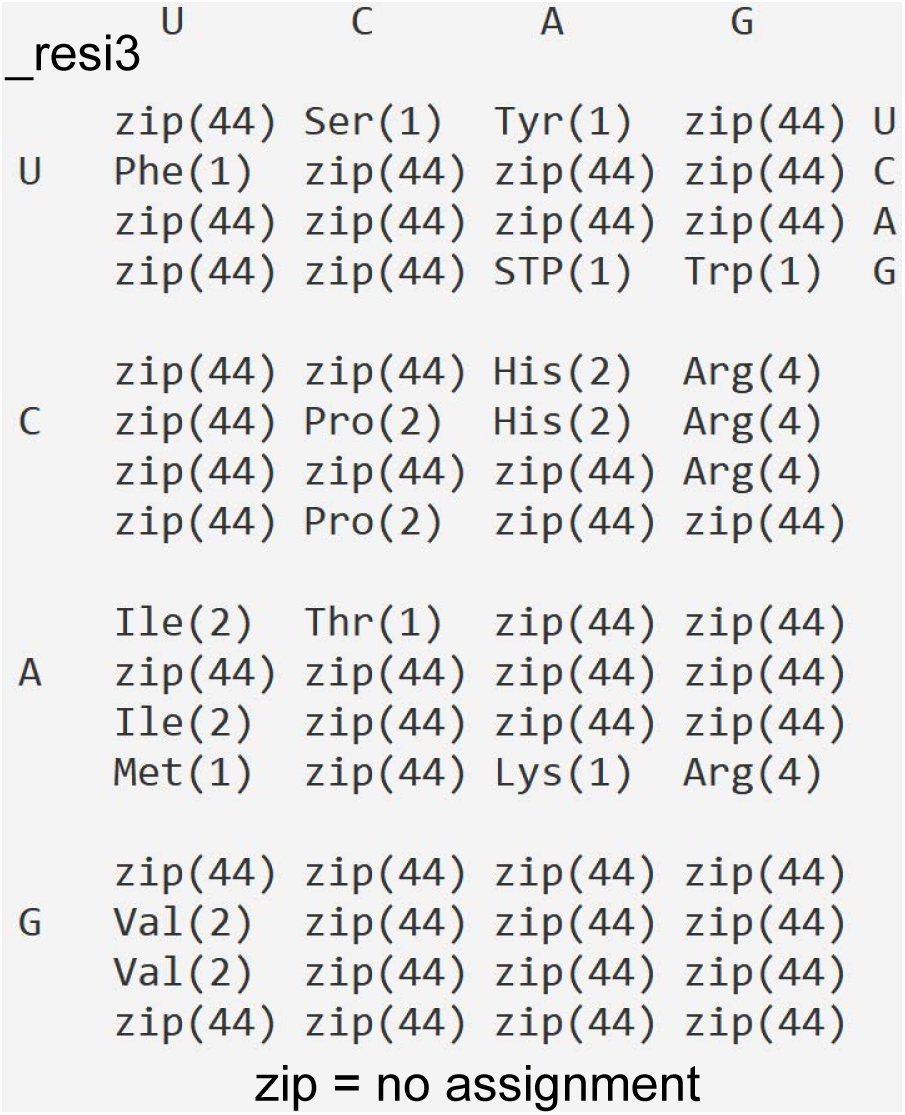
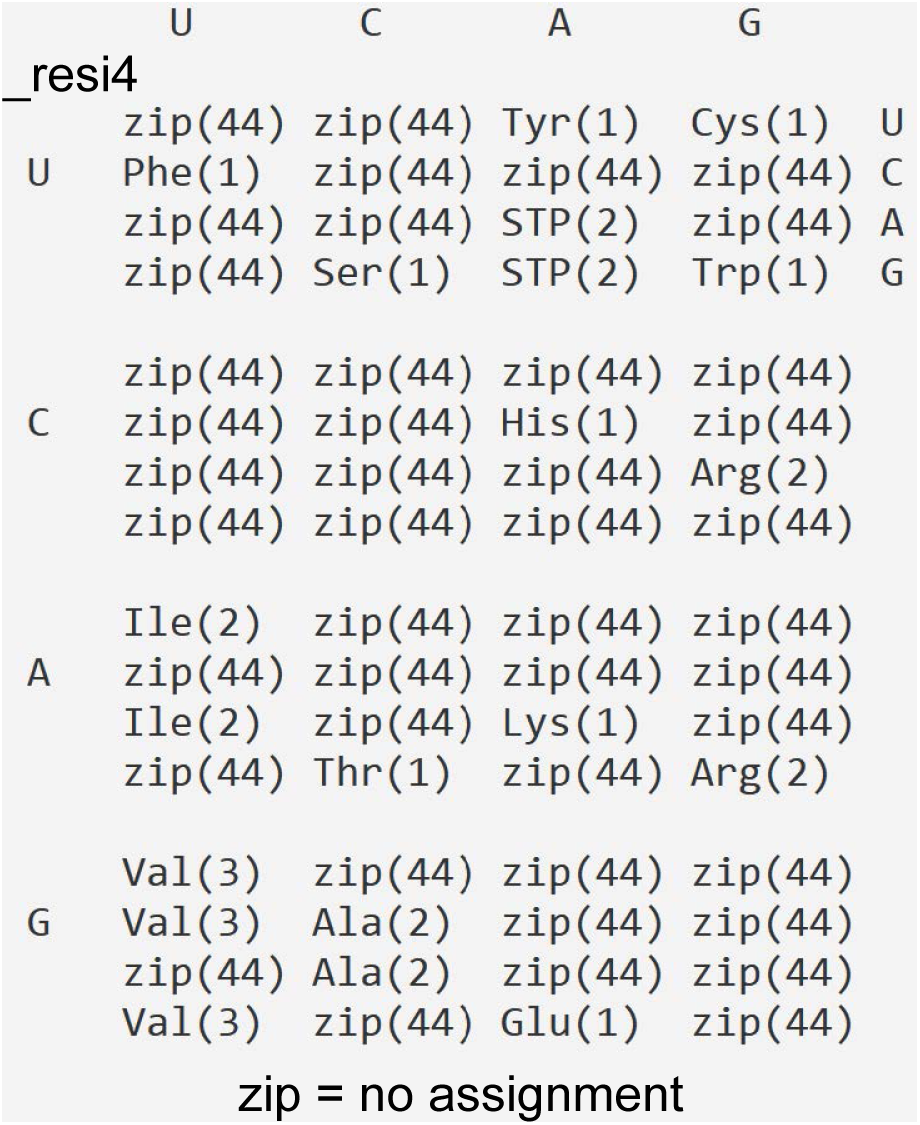
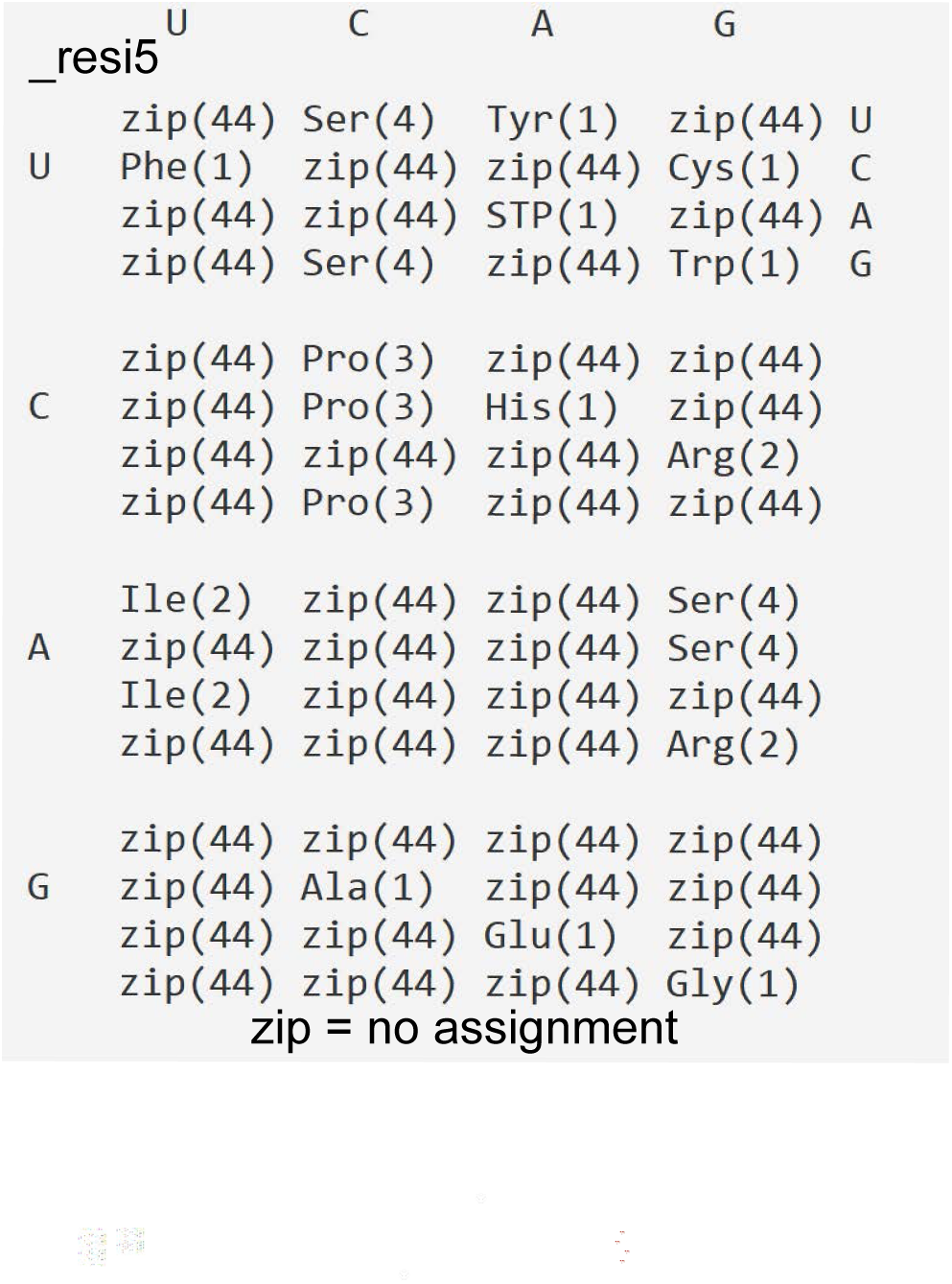

